# Understanding the effects of contact structures and information sharing on the FMD transmission among beef cattle farms

**DOI:** 10.1101/2020.04.27.063735

**Authors:** Chunlin Yi, Qihui Yang, Caterina M. Scoglio

**Author notes:** These authors contributed equally to this work (Chunlin Yi, Qihui Yang).

## Abstract

Moving infected animals and sharing contaminated vehicles are considered as the most potent ways for between-farm disease transmission. The objective of this study is to develop a network-based simulation model to investigate the effects of direct contact, indirect contact, and their combination on a hypothetical foot-and-mouth disease spreading between beef-cattle farms in southwest Kansas, US, and explore the effect of different types of information-sharing networks on preventing the disease spreading. Based on synthetic cattle and truck movement data in southwest Kansas, we build a farm-level contact network with three layers, a cattle movement layer (direct contact), a truck movement layer (indirect contact), and an information-sharing layer. Through scenario analyses, we compare the disease transmission dynamics, the distribution of outbreak epidemic size, and the disease breakout percentage of different contact structures – only direct contact, only indirect contact, and their combination. In addition, we evaluate different types of information sharing methods by comparing the epidemic size and the estimated economic loss. Simulation results show that neither direct contact nor indirect contact individually can result in a massive outbreak of the disease, but their combination plays a significant role. Additionally, we detect different probabilities of disease outbreaks by starting the simulations at different farms; starting at some farms with high capacity increases the probability of disease outbreaks. Three different information sharing-networks are developed and found effective in preventing the disease from spreading and reducing the economic loss. The information-sharing layer based on trading records has the best performance when compared with a random network and a geographic network.

## Introduction

Livestock diseases such as foot-and-mouth disease (FMD) are highly contagious and can lead to destructive economic losses in vast geography [1–4]. In epidemiological modeling, between-farm transmission via animal movement has been extensively studied and are often categorized as “direct contacts” [5,6]. In recent years especially after the 2001 FMD outbreak in the U.K, the between-farm transmission route of “indirect contact” which is through fomites such as personnel, vehicles, and agriculture tools, has drawn scholars’ attention [7–10]. For example, Thakur et al. [11] simulated the spread of porcine reproductive and respiratory syndrome virus among swine farms, and concluded that scenarios including only direct contact resulted in smaller epidemic sizes, compared with scenarios including both direct contact and indirect contact. In addition, the epidemic size under a scale-free network was larger than that under random and small-world networks. Rossi et al. built contact networks based on dairy cattle and veterinarian movement data, and showed that the direct contact and indirect contacts attributed to between-farm transmission from local and larger spatial scale, respectively [12]. Focusing on the same dairy farm system, Rossi et al. [7] further analyzed the impact of indirect contact route on FMD spreading and suggested that indirect contact has a larger impact on disease transmission compared to the direct contact network. As in many previous studies, they took each farm as a discrete single unite and used a simple Susceptible-Infectious-Susceptible compartmental model. Yang et al. [13] evaluated the effects of indirect contact and information sharing on FMD spreading. With each cattle agent simulated as the epidemiological unit and each truck following an independent clean-infected-clean cycle, they concluded that including indirect contact routes could significantly increase the epidemic size. In addition, they highlighted the effectiveness of mitigation strategies informed by information sharing, assuming that all producers participate in the information-sharing process. However, Yang et al. [13] did not explore the impact of partial participation in information-sharing on FMD transmission, which is one of the goal of this study.

In this paper, we will model the spread of an FMD-like disease among cattle farms focusing on the beef cattle industry in southwest Kansas (SW KS), United States. Accounting for $78.2 billion in cash receipts in 2015, the beef industry is an important sector in the U.S. economy. FMD outbreaks once occurred, mitigation strategies such as movement restrictions, culling animals on detected farms were often implemented, and such measures would almost result in devastating economic consequences. Several studies have estimated the potential economic impacts resulted from hypothetical FMD outbreaks in the United States. Pendell et al. estimated that financial losses of an outbreak constrained to Kansas would vary from $43 to $706 million, determined by the livestock species initially infected [14]. Schroeder et al. reported that the loss could be as many as 7,000–8,000 livestock herds and 16–18 million animals for a regional central U.S. outbreak [15]. Pendell et al. estimated that there would be around $16– 140 billion losses for an outbreak in the Midwestern U.S. originating from the proposed National Bio- and Agri-Defense Facility in Kansas [16].

Nonetheless, mitigation strategies informed by information sharing become increasingly promising thanks to the rapidly developed communication technology. Cattle production is a food supply chain system, and an increasing number of scholars have been studying the usage of information sharing as a method to improve supply chain resilience to disruptions [17,18]. Many studies have shown how information sharing awareness can effectively control the epidemic using multilayer modeling [19–21]. Similarly, we assume cattle farm operators equipped with information-sharing infrastructure will implement preventive measures after they are informed by their information-sharing partners even before animals show symptoms. As a result, disease spreading can be suppressed faster than traditional mitigation strategies. Farmers’ willingness to adopt the information-sharing process depends on a number of complicated factors such as risk attitudes, privacy, and transparency issues [22,23]. Many efforts are still needed to investigate those factors influencing farmers’ engagement and how the participation rate will affect the effectiveness of information sharing mitigation strategies.

Understanding the impact of direct contact and indirect contact on the epidemic dynamics is of great necessity to enhance the cattle production resilience. Therefore, this paper first investigates the potential impact of direct and indirect contact on FMD spreading among cattle farms in SW KS. We will generate a farm-level, weekly-resolution temporal network considering both direct and indirect contact, using the synthetic cattle movement and truck movement data generated in Yang et al. [24]. The simulation model will be built based on the Generalized Epidemic Modelling Framework (GEMF), which was developed by the Network Science and Engineering group at Kansas State University. Under different scenarios, extensive simulations will be carried out to test which type of contact structure has more impact on FMD spreading and how to develop better strategies to prevent livestock from infectious disease. The paper’s main objective is to assess how the information-sharing network and the fraction of participated farms can affect the epidemic size and estimated economic loss. The results are expected to shed light on the future dissemination strategies about the information-sharing infrastructure and used by relevant stakeholders.

Contributions of this paper can be summarized as follows:

1. Propose a model for the beef-cattle production system as a three-layer network: direct contact, indirect contact, and information sharing
2. Generate networks based on realistic cattle and truck movement data
3. Assess the role of each layer and its impact on disease transmission
4. Shed light on how the participation rate affects the effectiveness of information sharing

## Materials and methods

### Generation of the 3-layer network

We first generate a two-layer contact network based on the data from Yang et al. 2019 [24] where the cattle trading records and cattle transportation information (like the truck identifier, origin farm identifier, destination farm identifier, headcount of cattle carried by truck and transport date) among 18 ranches, 50 stockers, 233 feedlots, 4 packers, and 12 entry points in SW KS are represented. Here, we consider cattle and truck movements among ranches, stockers, and feedlots because the transmission between these farms are the most critical. Then, three types of information-sharing networks are developed as the third layer of the disease transmission network.

Since cattle movement from ranches to stockers in SW KS mainly occurs in the second half of the year, data for the last 26 weeks is used. To capture the dynamic characteristics of the contact network, weekly snapshot dynamic two-layer networks are generated and switched with the simulation calendar. In the first layer, the cattle movements are represented as directed weighted links with weights equal to the average number of cattle transported between premises per day in a week. In the second layer, similarly, undirected weighted links represent the average number of trucks commuting between two farms per day in a week. In the third layer, undirected and unweighted links are assigned between two farms if they are sharing information, which means they will get alerted if the other one is detected infectious.

### Individual-based epidemic model

To simulate the disease spreading between beef-cattle farms, we develop a farm-level, stochastic, individual-based epidemic model on GEMF, which can be used to simulate spreading processes on multilayer networks [25–27]. GEMF can numerically simulate any stochastic GEMF-based model, and is available in MATLAB, R, Python, and C programming language. In this work, we further adapt GEMF toolbox in Matlab for the simulations incorporating weekly snapshots of multilayer dynamic networks.

To explore the impact of the information-sharing network on preventing disease spreading, we introduce a new compartment quarantined (Q) to the traditional susceptible-exposed-infectious-removed (SEIR) model in this study to describe the epidemiology of FMD. In this transition process, each node can only be in one of the five compartment states; a susceptible node can become exposed if it is surrounded by infectious nodes in direct or indirect contact networks, the transition rate from S to E is 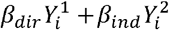, where *β*_*dir*_ and 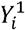 denote the infection rate and the number of contacts for direct contact, *β*_*ind*_ and 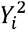 denote the ones for indirect contact. An exposed node is not yet infectious, but it will transit to the infectious state with rate λ. Finally, an infectious node will be removed with a removing rate δ. The farms can become quarantined with rate 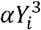 if they are connected to infectious farms in the information-sharing network.

The parameters used in this model are specified as follows: the infection rate for direct contact *β*_*dir*_ = 0.95*day*^−1^*calf*^−1^ [28], λ = 1/3.59 *day*^−1^ [29] which means the average incubation time is 3.59 days, *α* = 1*day*^−1^. For our base scenario, we use infection rate of indirect contact *β*_*ind*_ = 0.1*day*^−1^*calf*^−1^ and removing rate 8 = 1/14 *day*^−1^ [30].

In the network presentation, each node is a farm, and each link represents a movement contact between farms. The trade flow between different types of farms is unidirectional. For example, ranches sell calves to stockers but will not buy cows from other farms. Therefore, the disease can only spread along with the trade flow, and the network of cattle movement is directed. On the other hand, because the truck movement is assumed to be a round trip, the truck movement network is undirected.

For the SQEIR model, GEMF gives the individual-level Markov process for node *i* as

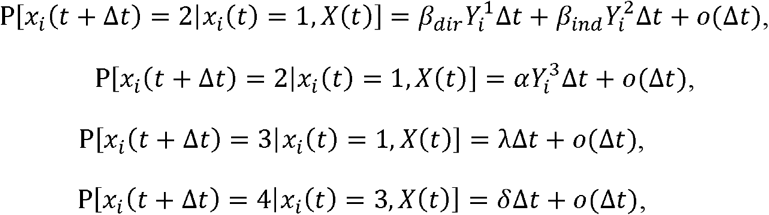

where *x*_*i*_ = 1, 2, 3, 4 *and* 5 correspond to the S, Q, E, I, and R state of node *i*, respectively. The value *X*(t) is the joint state of all nodes — the network state — at time *t*.

### Scenario Analysis

We designed several sets of simulations to evaluate the potential risk of FMD spread in the two-layer contact network and the effect of the information-sharing network on preventing disease spreading. We simulate 500 iterations of each scenario for 26 weeks during the second half of the year. We use the parameter values in the base scenario.

First, we simulate FMD transmission under 3 scenarios, namely by direct contact only, indirect contact only, and by direct and indirect contact. FMD virus is seeded in a randomly selected ranch in a randomly selected week in the first month.

Then, based on the farm capacity – the average number of cattle in this farm, we select the highest one in each production type as the high-risk farms. Then, we initialize the disease from each of the high-risk farms and compare the results with the scenarios where a random farm of each production type is seeded with the disease. The two-layer contact network with the combination of direct and indirect contact is used in these scenarios. The features of the disease are compared.

Additionally, sensitivity analysis for two key parameters -- the infection rate of indirect contact *β*_*ind*_ and the removing rate *δ* is performed for the 2-layers contact networks.

Lastly, we develop an information-sharing network as the third layer. We design 3 different types of information-sharing networks. The first one is a random network, the second one is based on the trade records in the past week, and the third one is based on the geographic distance between each farm.

The outcome of each simulation run is the total number of farms in each disease compartment over the simulation time. If there are at least 3 farms (1% of the total number of farms) infected at the end of the simulation time, we consider the disease breaks out, and otherwise, the disease dies out. From the outcomes, we extract the following measurements: 1) disease transmission dynamics, defined as the average fraction of farms in each compartment over time; 2) disease outbreak percentage, defined as the percentage of total simulations where the disease breaks out; 3) outbreak epidemic size, defined as the percentage of removed farms in iterations where the disease breaks out; 4) estimated economic loss.

## Results and discussion

### Effect of contact structures

The results of the average epidemic size over simulation time and distribution of epidemic size for different contact structures are presented in Fig. 1.

**Figure 1.**
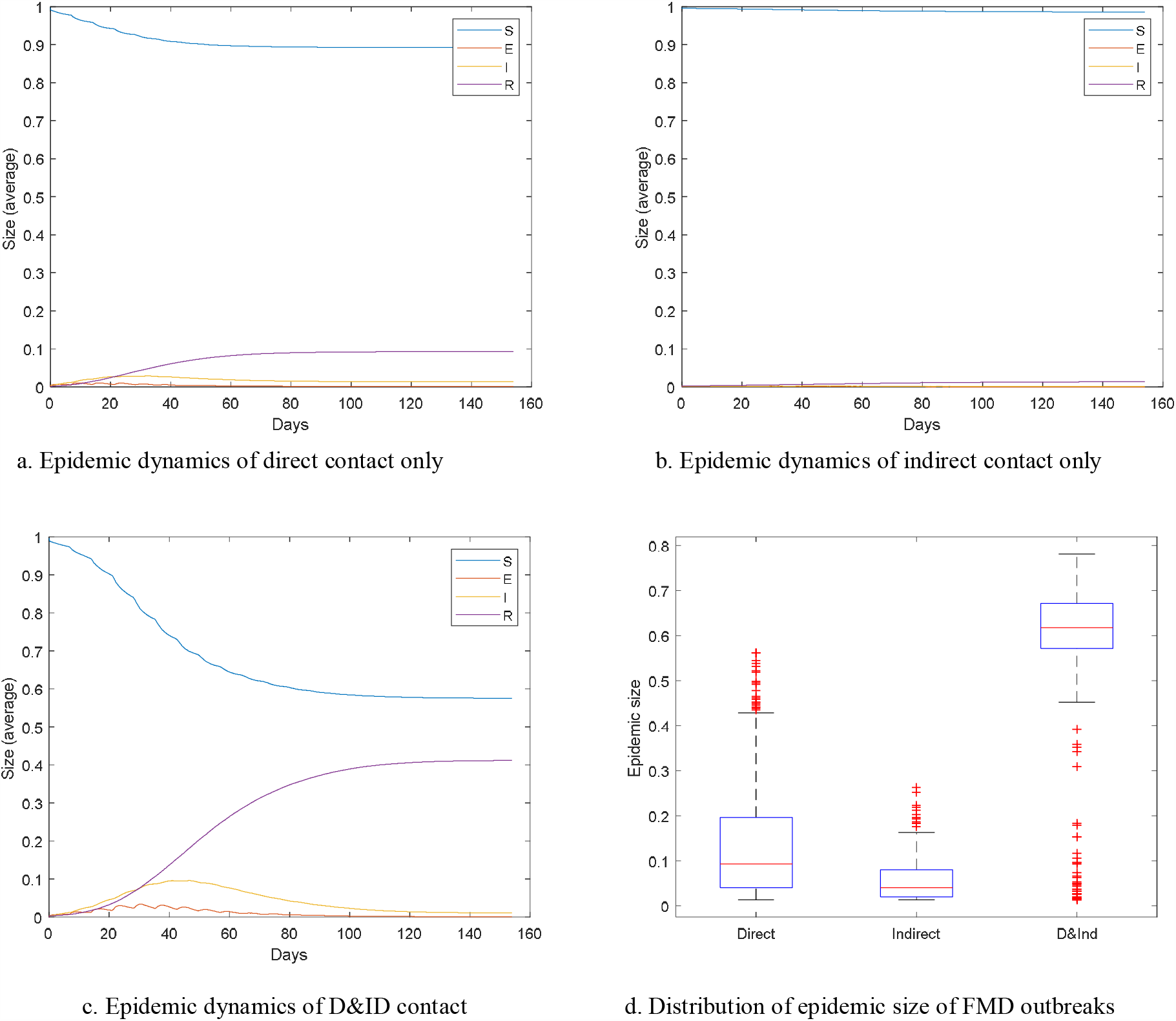
Disease transmission dynamics and distribution of outbreak epidemic size for the direct-contact-only network, indirect-contact-only network, and the combination of the direct and indirect contact network.

The disease outbreak percentages of these scenarios are 67% (335/500) for direct contact only network, 19.4% (97/500) for indirect contact only network, and 71.2% (356/500) for direct and indirect contact network. Fig. 1 (a) ∼ (c) show the disease transmission dynamics after introducing the FMD virus to a random ranch. The direct contact scenario (a): compartment R reaches the steady-state 0.092 at 60th day; the indirect contact scenario (b): the disease can hardly spread; direct & indirect contact scenario (c): compartment R arrives at steady state 0.401 at around the 100th day. Fig. 1 (d) reflects the simulation results where the disease spreads at least to 3 farms. For direct contact only network, the median of outbreak epidemic size is 0.093; for indirect only contact network, outbreak epidemic size in most cases is less than 0.1. As for direct and indirect contact network (D&ID), the median of outbreak epidemic size is 0.632, which is much larger than the scenario with only one contact layer.

Comparing the results from Fig. 1, we find that disease can hardly spread through a single contact network. However, when we combine these two layers, we see dramatic growth in the epidemic size and endemic duration. In this study, the direct contact network is an acyclic directed network, which prevents the reverse transmission of the disease. Even though the indirect network is undirected, the infectious rate for this layer is quite small, which prohibits the disease spread. However, the network with direct and indirect contact combines the contributions of the two transmission mechanisms to the spreading and generates disease outbreaks.

### Effect of infecting the high-risk farms of different production type

The following scenarios are designed to explore the effect of infecting the high-risk farms on disease spread.

1. We initialize the disease from the high-risk ranch (HRR), the ranch with the highest capacity, at a random week in the first month. From Fig. 2 (a), the time to reach a steady state is at around 100 days, and the average epidemic size is 0.559, larger than the case with random selected infectious ranch, shown in Fig. 2 (b), which means that the overall risk would be higher if ranch with the highest capacity is initially infected. There are 483 occurrences of the secondary case over 500 iterations when the FMD virus is introduced in HRR. Namely, if the ranch is initially infected, there will be more than 90% probability that the disease will break out. However, the probability of a disease outbreak when the virus is introduced from a randomly selected ranch is around 70% (356/500). On the condition that the disease broke out, the distribution of epidemic size for both scenarios are quite similar, as is shown in Fig. 2 (b). Therefore, if the disease is started from the HRR, there will be a higher probability that the disease will break out, but the epidemic size of the FMD outbreak will hardly be influenced.
2. We initialize the disease from the high-risk stocker (HRS) at a random week in the first month and compare it with initializing the disease from a random stocker. Fig. 3 (a) shows the disease transmission dynamics when the HRS is initially infected; the steady state of R is 0.245; Fig 3. (b) shows the disease transmission dynamics when a random stocker is initially infected; the steady state of R is 0.129. Distribution of outbreak epidemic size is shown in Fig. 3 (c); the medians of the outbreak epidemic size for the cases where the HRS and a random stocker is infected are 0.568, 0.601. Among 500 simulations for each scenario, there are 271 disease outbreak simulations when the HRS is initially infected; 187 simulations where the disease breaks out when a randomly selected stocker is infected, which shows that if the disease is initialized from the high-risk stocker, the probability for disease outbreak is higher than that in the case where disease starts from a randomly selected stocker. Comparing the results in Fig. 2 and Fig. 3, we notice that the difference between the disease outbreak percentages is large. Disease starting from ranches has a higher probability of spreading. Furthermore, a disease initialized from a ranch has a more concentrated distribution and greater value of outbreak epidemic size than the case where the disease starts from a stocker. The main reason for these is that ranches can reach more farms in a direct contact network than stockers. Therefore, the risk of an outbreak when initializing the disease from a ranch is greater than that when starting from a stocker.
3. When initializing the disease from a high-risk feedlot or a randomly selected feedlot, we find that the disease cannot spread because the feedlot cannot reach any other farms in the direct contact network.

**Figure 2.**
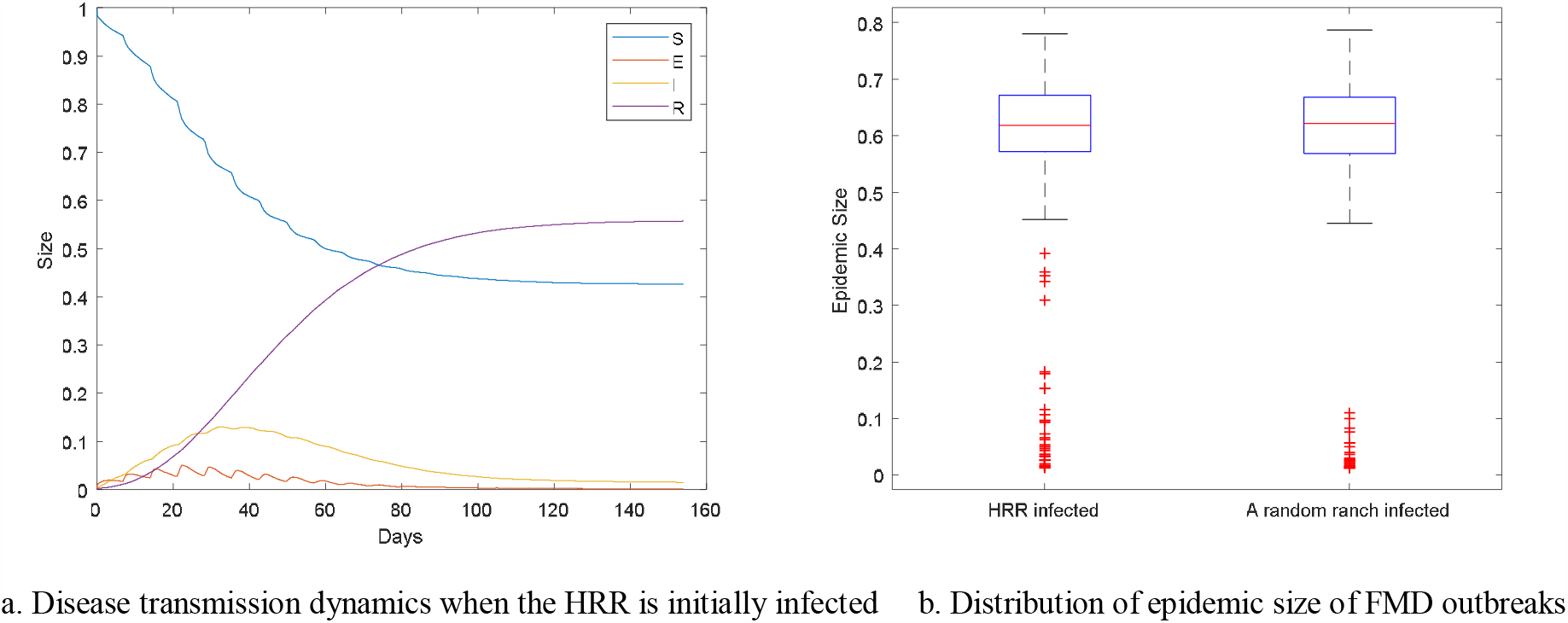
Disease transmission dynamics and distribution of epidemic size when the ranch with the highest capacity is initially infected.

**Figure 3.**
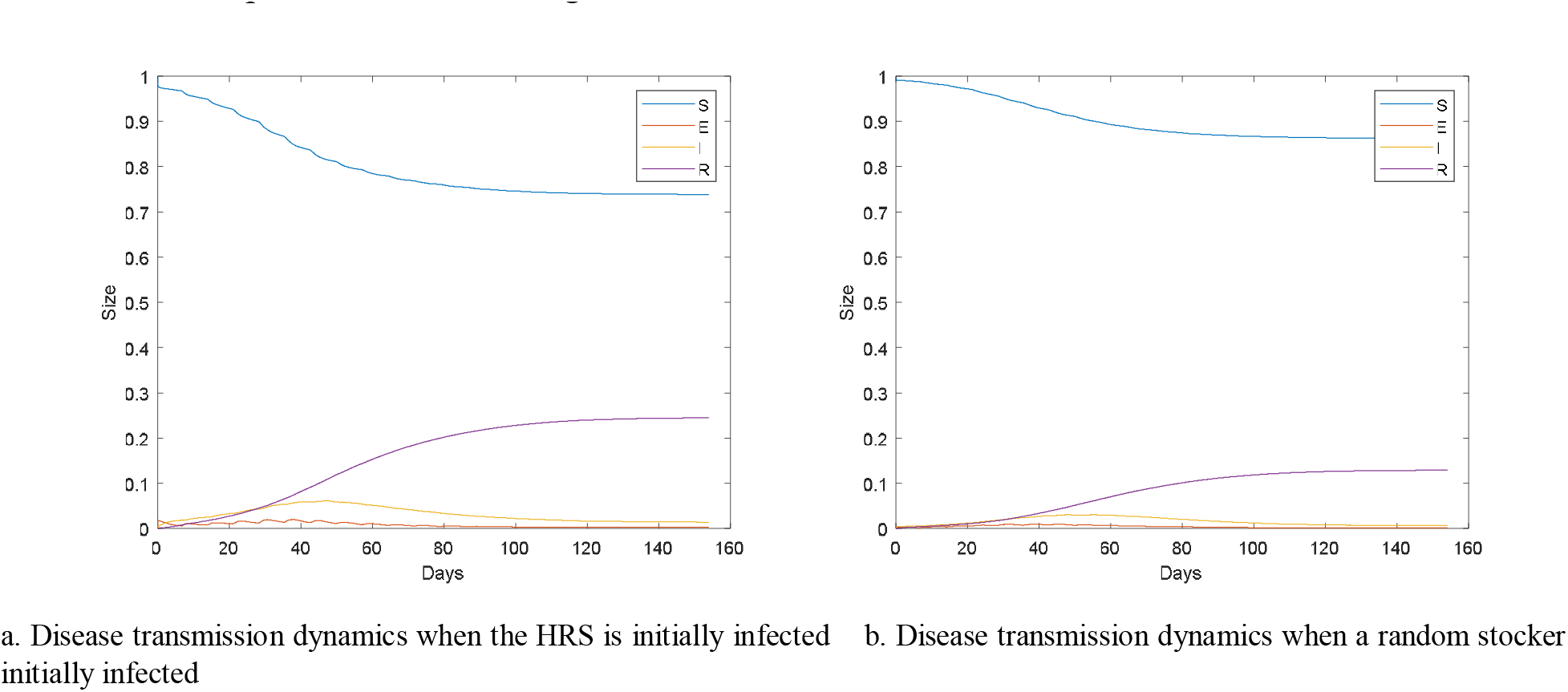
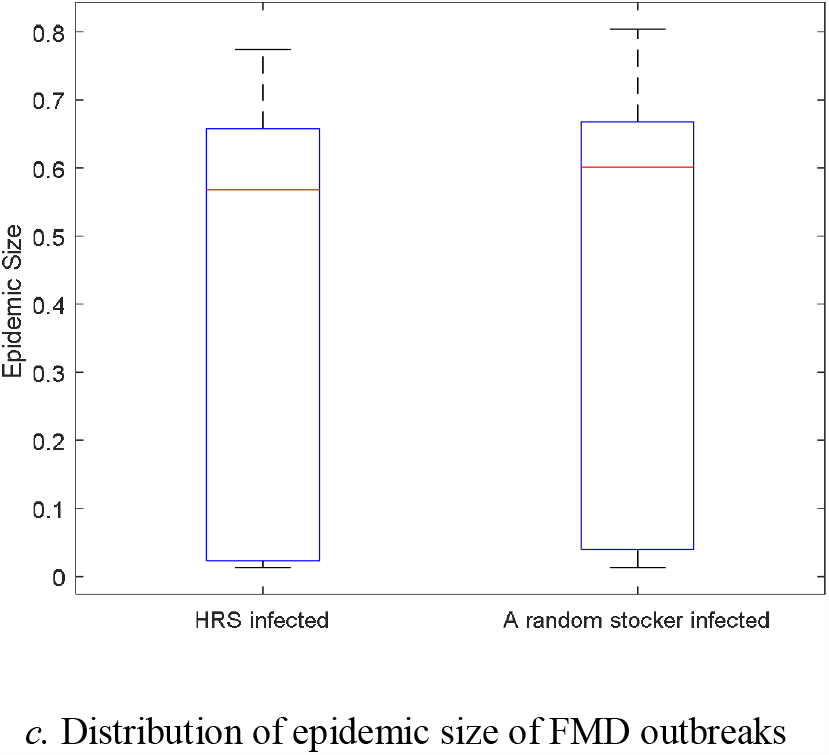
Disease transmission dynamics and distribution of outbreak epidemic size when the HRS and a randomly selected stocker is initially infected, respectively.

### Sensitivity analysis on removing rate δ and indirect infection rate *β*_*ind*_

We conduct a sensitivity analysis through a two-part approach. In the first part, we study the effect of the removing rate *δ* on FMD transmission. The reciprocal of the removing rate indicates the time needed by the regional administration to take actions from the day when the first case of the disease is detected. Therefore, how to choose this parameter is of great importance for disease control. We vary the removing rate from 1/4 to 1/23 while keeping other parameters in the base scenario unchanged. We perform 500 simulations for each *δ*. The fraction of infected farms over the total number of farms in southwest Kansas is recorded for each value of 8. The outbreak epidemic size distribution and the outbreak percentage are shown in Fig 4.

**Figure 4.**
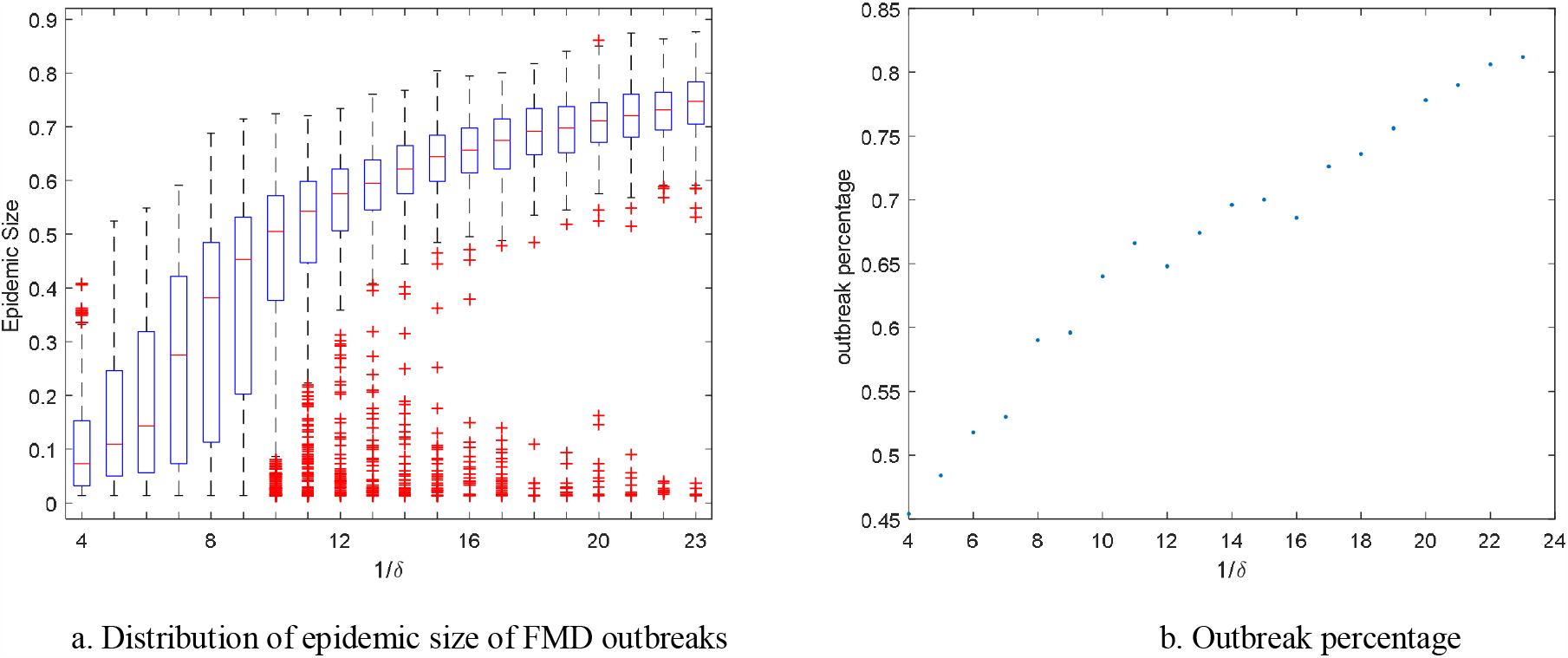
Distribution of epidemic size and outbreak percentage when *1/δ* varies.

In this section, we study the effect of removing rate on FMD transmission. 1 over the removing rate indicates the time needed by the regional administration to take action, from the time the first case of the disease was reported. Therefore, how to choose this parameter reasonably is of great importance for disease control. In total, there are 20 scenarios in this simulation, with 500 iterations of simulation for each scenario. The data distribution is shown in Fig. 4 (a), and the die-out percentage is shown in Fig. 4 (b).

Fig. 4 (a) shows that the outbreak epidemic size increases when the removing rate 8 decreases. From Fig. 4 (b), we could see that the outbreak percentage is strongly correlated with 1/*δ* with the linear correlation coefficient 0.978.

In the second part, the effect of changes in the indirect infection rate *β_ind_* on disease transmission is studied. The indirect infection rate reflects the ability of the disease vector to spread pathogenic bacteria. Adjusting the value of this parameter is equivalent to taking certain measures, such as cleaning and disinfecting vehicles. In this set of simulations, we have selected 20 different values ranging from 0.01 to 0.20 while keeping other parameters in the base scenario unchanged. Simulations results are presented in Fig. 5.

**Figure 5.**
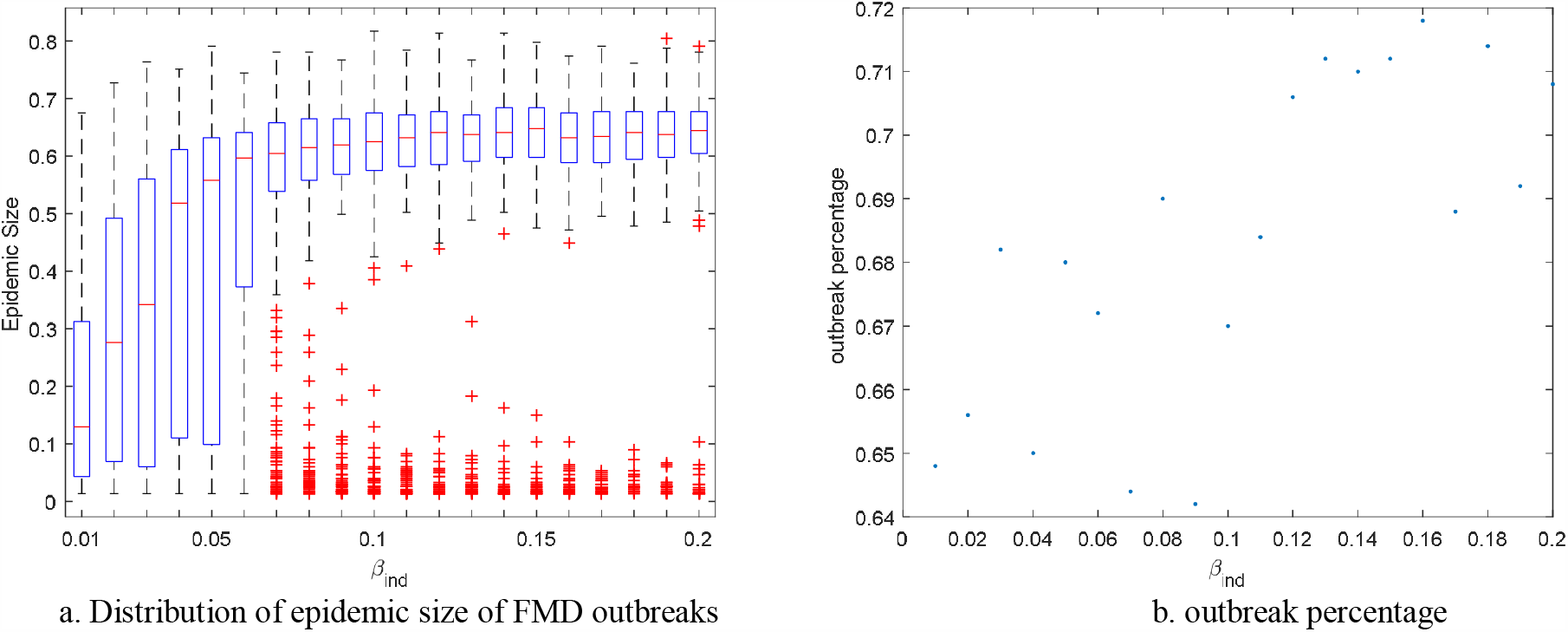
Distribution of epidemic size and outbreak percentage when *β*_*ind*_varies.

As shown in Fig. 5 (a), the medians of epidemic size increase very quickly when *β_ind_* is below 0.07. However when *β_ind_* >0.07, there is little difference in the median of epidemic size. Therefore, we can conclude that by controlling the indirect infection rate *β_ind_* can significantly reduce the overall number of infected farms. In addition, it is obvious that the epidemic size has a saturating effect on the parameter *β_ind_*. In other words, the epidemic size (median and quartiles) changes slightly with the increase of the parameter once it reaches a certain value 0.06. When beta is between 0.01 and 0.06, the epidemic size grows rapidly. Once *β_ind_* is above 0.06, the epidemic size reaches the peak, with a median value at around 0.63. However, from the previous Fig. 4 (a), the median epidemic size can reach 0.75, suggesting that the removal rate limits the further increase in epidemic size. In Fig. 5 (b), the distribution of data points is more scattered and the variance of outbreak percentage is 0.076, which is much smaller than the removal rate *δ* 0.358. Besides, the correlation coefficient between the indirect infection rate and the die-out percentage is 0.716, which means that the removing rate has a higher impact on the disease outbreak probability than the indirect infection rate.

### Effect of the information-sharing network

Susceptible farms can become quarantined when they receive information from other infected farms if they are connected in the information-sharing layer. Quarantined farms will stop business with the other farms so that they will not be infected. For simplicity, we assume that the quarantined farms will stay in the state till the epidemic ends. To evaluate the economic loss caused by the epidemic, we introduce a cost function *loss* = *Rx* + *Qy*, where *x* is the average loss for one removed farm, y is the average loss for one quarantined farm, *R* and *Q* are respectively the average number of removed farms and the average number of quarantined farms. Considering that infected farms will cull the cattle to clear the infection, and the quarantined farms have a higher cost to keep all the cattle, *x* should be greater than *y*. Take *y* as 1, and assume *x* equal to 2; we have the cost function *loss* = 2*R* + *Q*.

Three different types of information-sharing networks are developed and added as the third layer of the disease transmission network. Different scenarios of participation rates are designed for each network.

1. Random information-sharing network with random participating farms The first information-sharing network is an Erdos Renyi random network with probability *p* = 0.007 of creating a link between any pair of nodes, which is the density of the cattle movement network. We randomly included 10% to 100% participating farms of each production type and have the results shown in Fig. 6. In Fig. 6, the median of epidemic size decreases from 0.62 to 0.34 with more farms included, while the median of quarantine size increases from 0.01 to 0.4. The increased value of quarantine size is greater than the decreased size of infected farms. Even though quarantining farms leads to less economic loss than culling cattle if the farms are infected, the total loss of the epidemic is not strictly decreasing with more farms included, as shown in Fig. 6 (c). The minimum loss (0.64) is obtained when all farms are selected.
2. Random information-sharing network with highest-capacity participating farms We use the random network, but include the farms with the highest capacity. The results are shown in Fig. 7. In Fig. 7, the median of epidemic size decreases from 0.62 to 0.34 and the median of quarantine size increases from 0.01 to 0.41 as more farms are included in the information-sharing network, which is very similar to scenario 1). The minimum estimated economic loss is 0.66 when participation rate is 100%. Comparing with the 1) scenario, the epidemic size decreases faster while the distributions of quarantine size are very similar when the participation rate is small. The estimated economic loss changes slightly when more than 50% farms participate. The farms with the highest capacities usually have more business connections with other farms so that they are more contagious. Including partial high-capacity farms in a random information-sharing network can minimize the economic loss.
3. Trade-recording information-sharing network and highest-capacity participating farms The third one is developed based on the trade records during the last week. The farms having contacts with infectious farms will be quarantined with a probability. We include farms with the highest capacity of every production type and have the results shown in Fig. 8. The median of epidemic size decreases to 0.15. The quarantine size increases to 0.22 as more farms are included in the information-sharing network, and the minimum loss is 0.35 when 70% of farms are included, as shown in Fig. 8 (c). Based on the results, this information-sharing network has a much better performance on disease prevention and quarantine minimization. The optimal is quite less than that of the former two scenarios. This network is not random so that the transmission of the virus is traceable.
4. Geographic information-sharing network with all farm participating The last information-sharing network is developed based on the geographic distance between each farm. If the distance between two farms is shorter than a preset radius, those farms are connected in the information-sharing network. The distance between each pair of farms ranges from 0 to 18,748km. The epidemic dynamics are shown in Fig. 9. The epidemic size decreases very fast while the size of quarantined farms increases even faster as the distance enlarges. In other words, the geographic information-sharing network can make the epidemic size decrease to 0 but will have most farms in quarantine. Therefore, the total loss of the epidemic is still high, which results from too many quarantined farms. In Fig. 9 (c), the minimum estimated economic loss 0.58 is achieved when the distance is 1000km. We use this optimal geographic network to explore the effect of the participation rate of the highest capacity farms. In Fig. 10, the median of epidemic size decreases from 0.45 to 0.15. The quarantine size increases to 0.67 as more farms are included in the information-sharing network, and the minimum loss is 0.54 when 80% of farms are included. In this case, the participation rate is of little importance in reducing estimated economic loss.

**Figure 6.**
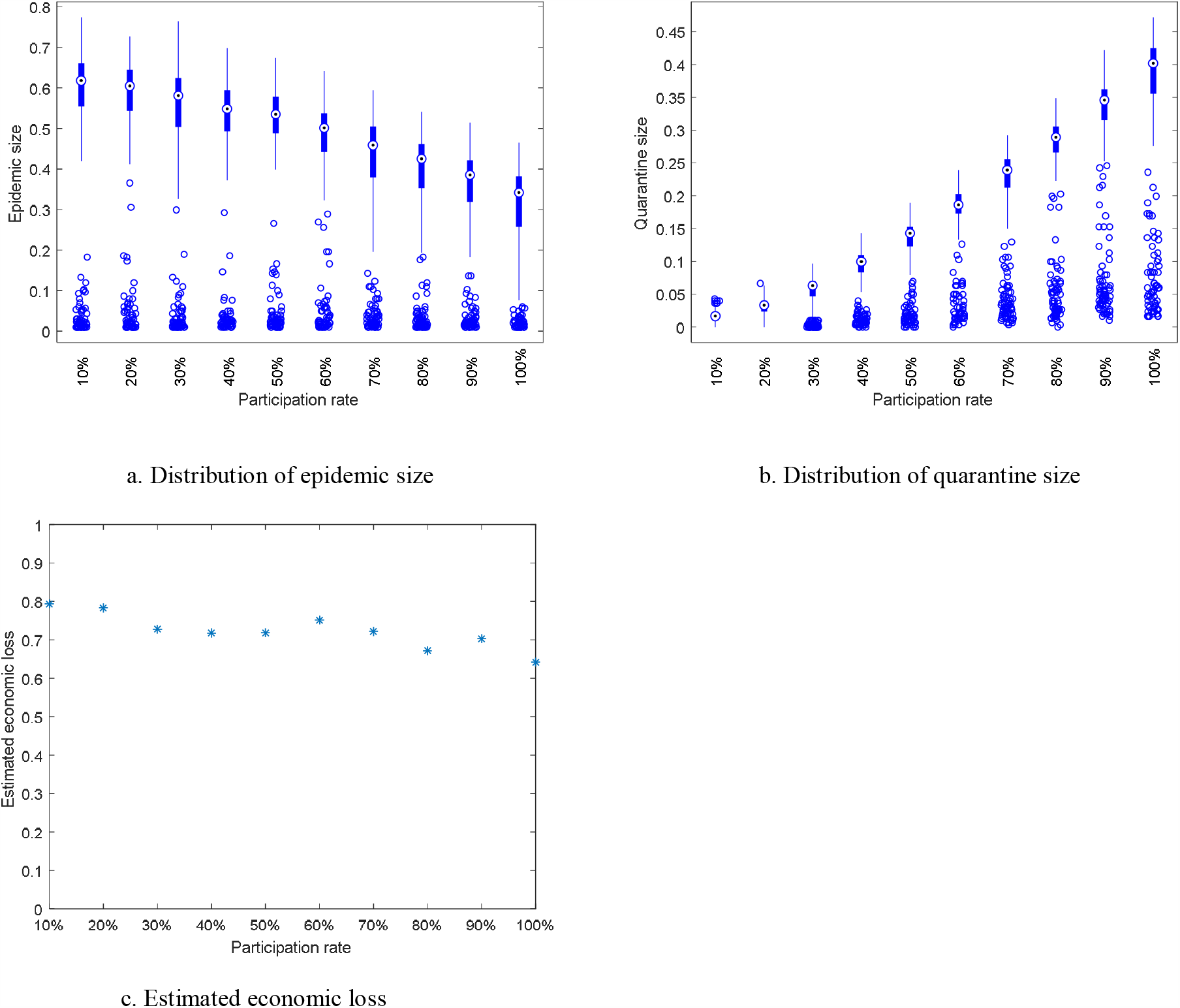
Distribution of epidemic size and quarantine size and estimated economic loss of random information-sharing network with random participating farms.

**Figure 7.**
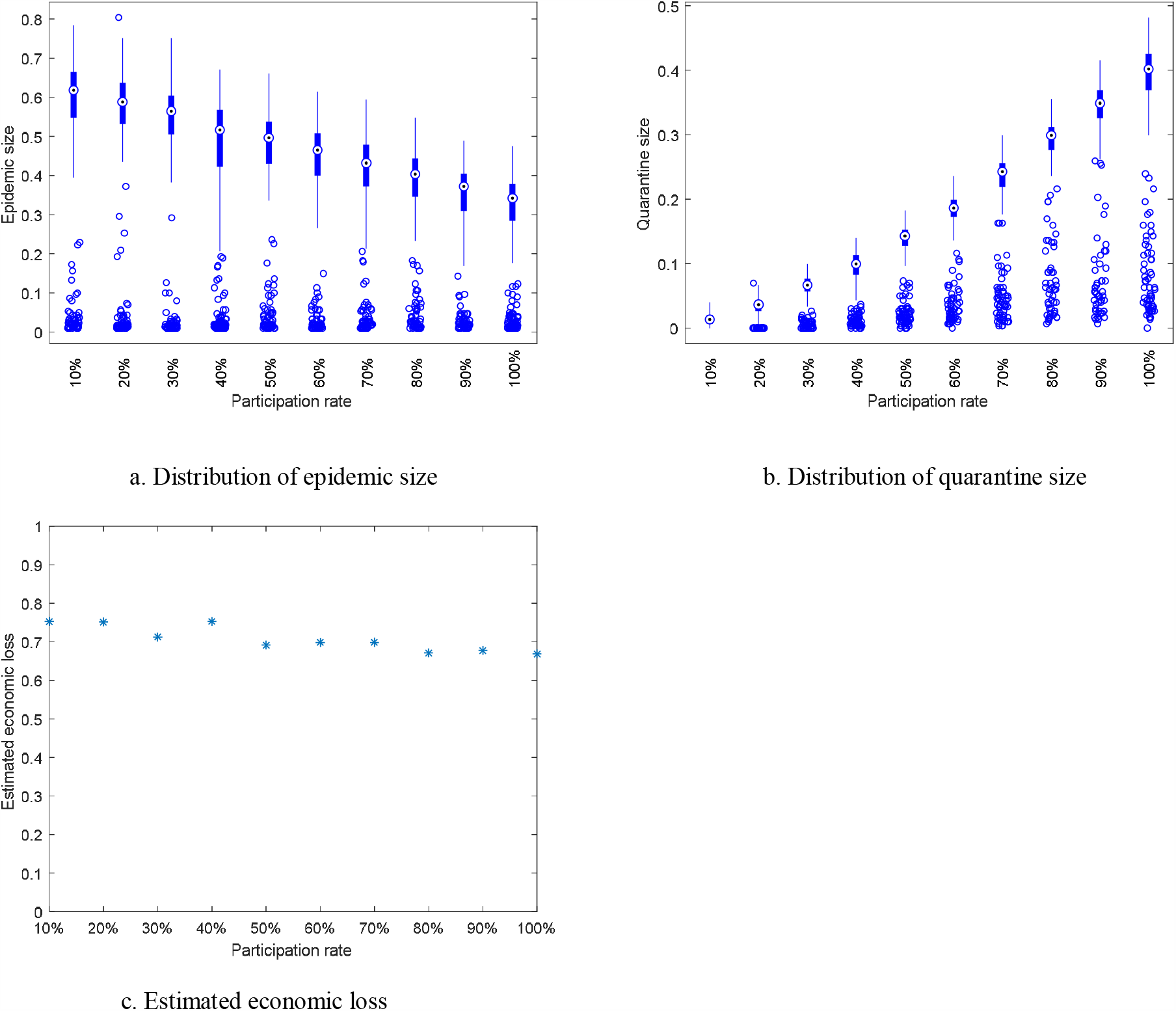
Distribution of epidemic size and quarantine size and estimated economic loss of the random information-sharing network with highest-capacity participating farms.

**Figure 8.**
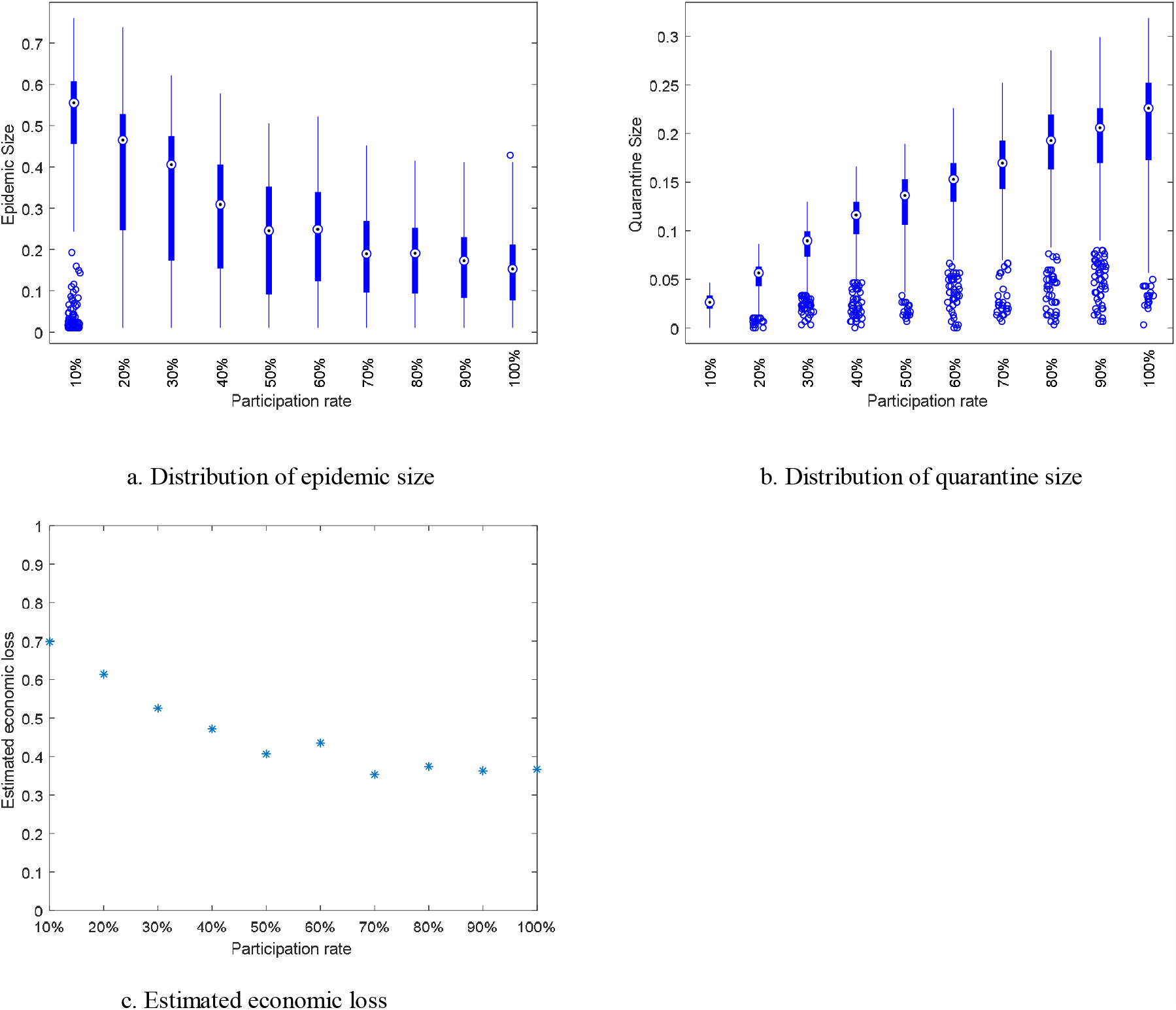
Distribution of epidemic size and quarantine size and estimated economic loss of trade-recording information-sharing network with highest-capacity participating farms.

**Figure 9.**
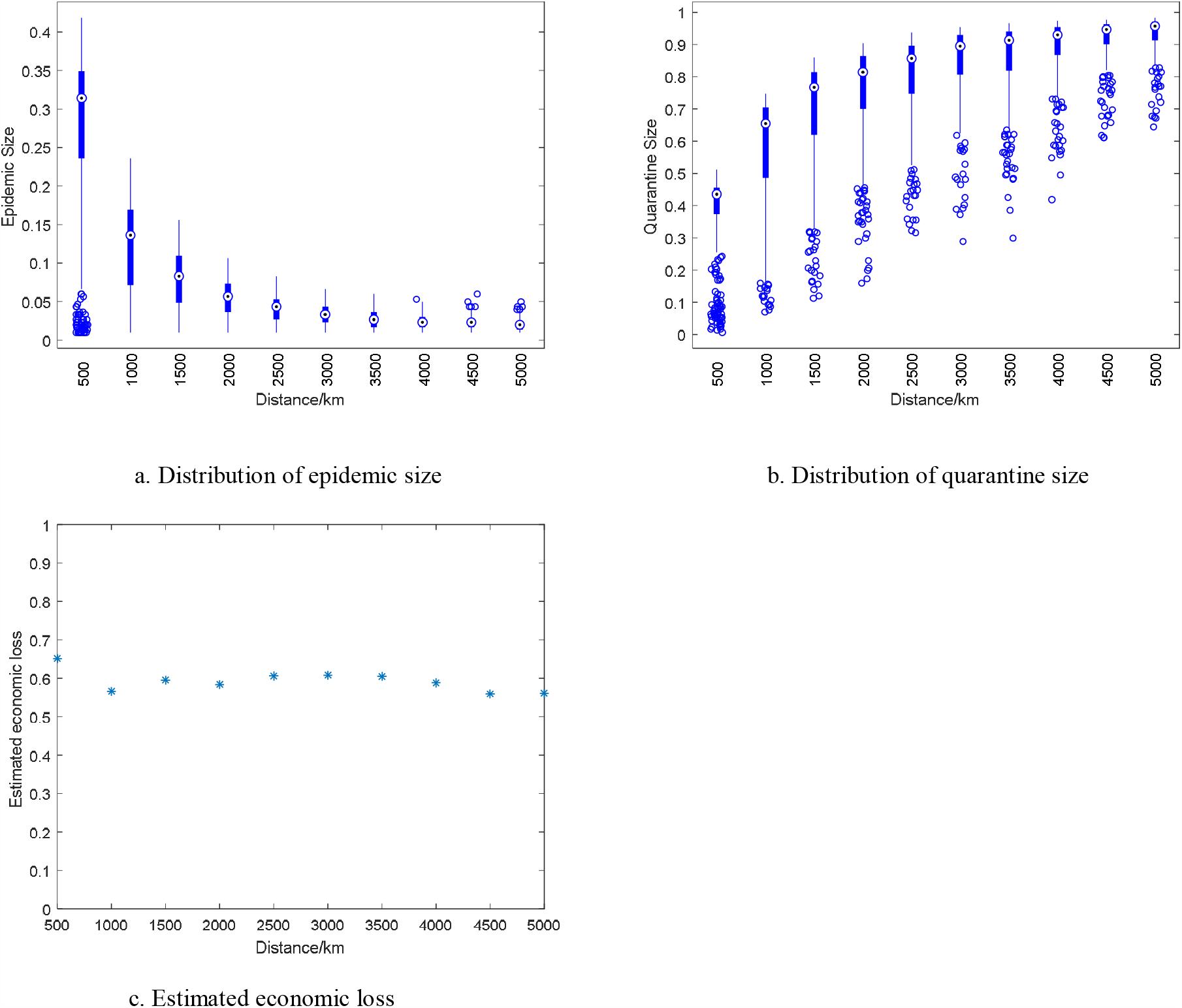
Distribution of epidemic size and quarantined size and estimated economic loss of the geographic information-sharing network.

**Figure 10.**
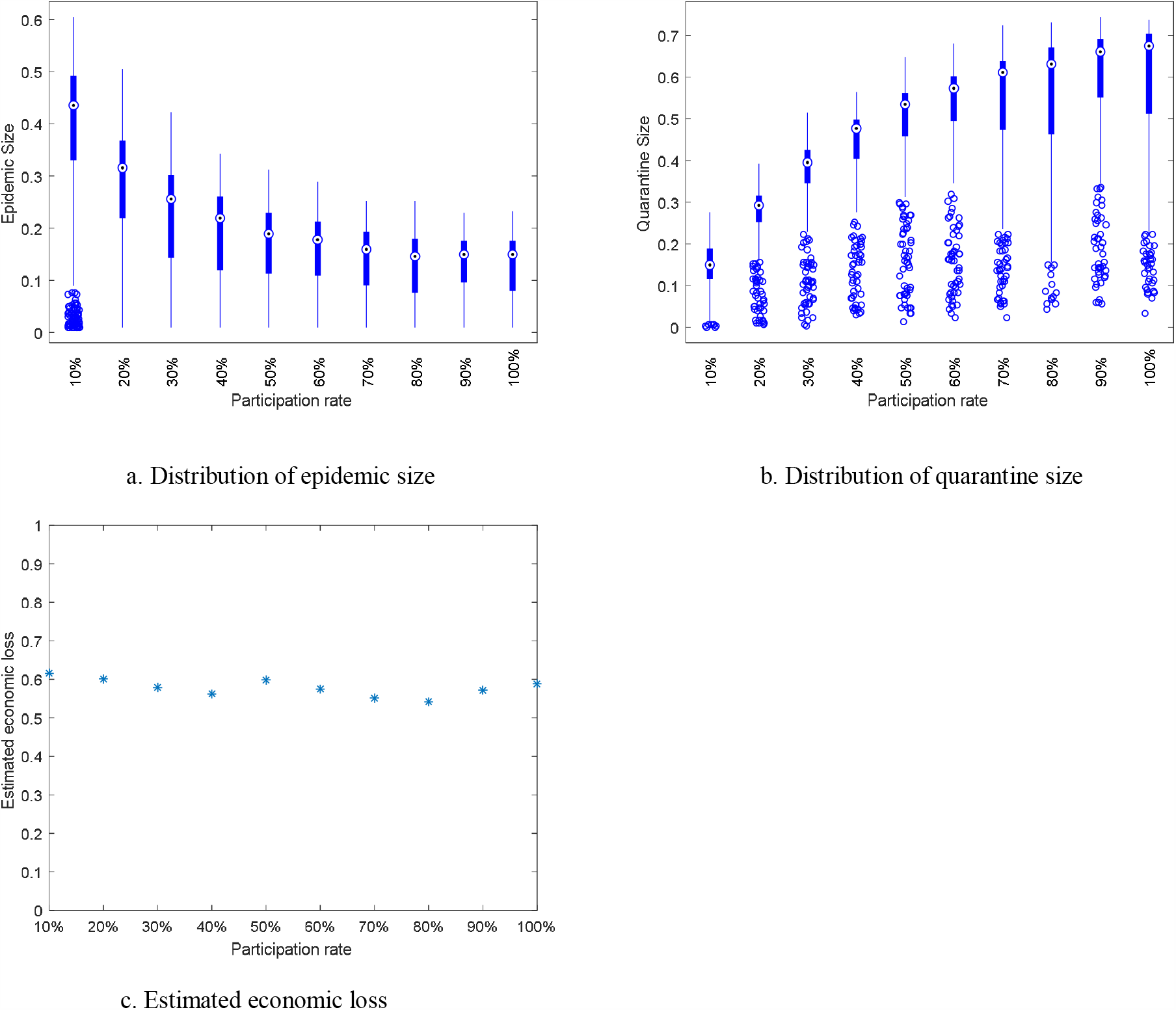
Distribution of epidemic size and quarantined size and estimated economic loss of geographic information-sharing network.

Comparing the results of all scenarios above, the trade-recording information-sharing network with the 70% highest capacity participating farm has the best performance on reducing the estimated economic loss. For each information-sharing network, the epidemic size is smaller with a higher participation rate. However, the estimated economic loss can be increasing with more farms included because of the loss from quarantining too many farms. Participating rates play an important role in reducing the estimated economic loss of the epidemic for the trade-recording information-sharing network. We don’t need to include all the farms into the trade-recording information-sharing network, but only part of the farms with the highest capacity to have the best results. Geographic information-sharing networks with a large connection radius can stop the disease from spreading with the price of quarantining almost all farms.

## Conclusions

The current study has developed an infectious disease spread model to evaluate the impact of the different contact structures on FMD outbreaks among farms in southwest Kansas, US. Based on a farm-level, weekly-resolution temporal network of direct contact and indirect contact, FMD spreading scenarios are simulated and analyzed by considering 1) only direct contact, 2) only indirect contact, 3) their combination. We verify that disease can hardly spread among farms on a single contact-type network. The direct contact influences the disease outbreak probability; the indirect contact can increase the epidemic size dramatically when incorporating with the direct contacts. The study further reveals that infecting ranches or high-risk farms can result in a larger disease outbreak. In addition, by studying the relations between disease outbreak percentage and epidemic size with removing rate and indirect infection rate, we find that the removing rate is highly related to the outbreak percentage so that the early detection and isolation of infected farms can greatly reduce the probability of epidemic outbreak and that the epidemic size increases very fast with the indirect infection rate when the infection rate is lower than 0.06 so cleaning the vehicles and other equipment shared with other farms frequently is an effective way to prevent FMD spreading. What’s more, three different information-sharing networks are added as the third layer to prevent the disease transmission and the effect of participation rate is explored. Trade-recoding information-sharing network has the best performance on minimizing the estimated economic loss among random information-sharing network and geographic information-sharing network. Additionally, we find that only including a certain part of farms with the highest capacity in the information-sharing network can achieve the best results.

Factors that we did not include in this study are that disease might spread through packers and the markets. In fact, the number of packers and trade markets is quite small compared to the number of farms in a specific area. In addition, cattle just stay in packers and markets for a short period of time. The model for disease transmission through packers and markets should be different. The cost function can be more complex and the quarantine duration is not considered.

## Acknowledgments

This work was supported by United States National Science Foundation under Grant Award CMMI-1744812. The funders had no role in study design, data collection and analysis, decision to publish, or preparation of the manuscript.

